# Simulation of cryo-EM ensembles from atomic models of molecules exhibiting continuous conformations

**DOI:** 10.1101/864116

**Authors:** Evan Seitz, Francisco Acosta-Reyes, Peter Schwander, Joachim Frank

## Abstract

Molecular machines visit a continuum of conformational states as they go through work cycles required for their metabolic functions. Single-molecule cryo-EM of suitable in vitro systems affords the ability to collect a large ensemble of projections depicting the continuum of structures and assign occupancies, or free energies, to the observed states. Through the use of machine learning and dimension reduction algorithms it is possible to determine a low-dimensional free energy landscape from such data, allowing the basis for molecular function to be elucidated. In the absence of ground truth data, testing and validation of such methods is quite difficult, however. In this work, we propose a workflow for generating simulated cryo-EM data from an atomic model subjected to conformational changes. As an example, an ensemble of structures and their multiple projections was created from heat shock protein Hsp90 with two defined conformational degrees of freedom. All scripts for reproducing this workflow are available online. ^1^

## Introduction

Molecular machines, macromolecular assemblies comprised of proteins or nucleoproteins that perform basic life functions, undergo numerous conformational changes and visit a large number of states in the course of their work cycles. For some molecular machines, such as the ribosome, it has been possible to develop *in vitro* systems with all components necessary to run productive work cycles delivering the product; in this instance a protein. Even though the atomistic composition of molecules ultimately defines a finite step size, these states form a quasi-continuum due to the complexity of the structures involved and the large number of steps in their interactions.

Through application of advanced imaging techniques, single-particle cryogenic electron microscopy (cryo-EM)^2–4^ allows macromolecules in an *in vitro* suspension to be experimentally visualized en masse after rapid freezing in vitreous ice at a rate that is assumed to be faster than the overall reconfiguration time of the system. ^5^ As a result, the ensemble of frozen molecules closely approximates the equilibrium distribution of states immediately prior to being frozen. Cryo-EM aims to capture the macromolecule in its complete form: encompassing all projection directions across its continuum of allowable states.

Recent advances in cryo-EM resolution have inspired the development of several data-analytic approaches that aim to systematically describe the characteristics of the ensemble. These methods rely on dimensionality reduction to gain insight and understanding of the most significant phenomena captured in the data. Linear dimensionality reduction methods have been proposed using principal component analysis (PCA) on the 3D molecular structures.^6–11^ However, these techniques are unreliable for mapping intrinsically nonlinear conformational spaces of molecular machines. To circumnavigate this problem, nonlinear techniques have also been applied^12–15^ via the Diffusion Maps method^16,17^ to represent the intrinsic conformational manifold of the cryo-EM data. This manifold can then be traversed to fully recover the molecular machine’s full set of most significant conformational coordinates. Recently, taking a page from both books, the Laplacian spectral volumes method^18^ combines both linear and nonlinear dimensionality techniques in its search of a lower-dimensional description of the data using only a few effective coordinates. A range of other techniques for analyzing cryo-EM continua are also being implemented,^19^ extending into work using deep neural networks.^20^

Regardless of technique, once the state space has been sampled and organized along these coordinates, the distribution of sightings at each state can be transformed through the Boltzmann relationship into a free-energy landscape. ^21^ The free-energy landscape provides the thermodynamic likelihood of a hop from any one state to any of its immediate neighbors. The topology of this landscape characterizes the system’s navigational potential, with deep wells representing distinct regions of conformational states and mountainous regions constraining the transitions between them. Further, since specific sequences of conformations give rise to biomolecular function, this descriptive mapping also accounts for the molecular machine’s functional dynamics.^22^ If these algorithms are successful, the potential exists for creating the conformational free-energy landscape for any molecular machine capable of being visualized by single-particle cryo-EM.

Since the energy landscapes of molecular machines are not experimentally known, these techniques cannot be immediately validated. Instead, they must first be tested on simulated cryo-EM images of a synthetically-designed ensemble having known structural and dynamical properties. To this end, we have constructed a workflow for creating a synthetic quasi-continuum of structures from an experimentally-relevant ground-truth model. The resulting dataset emulates the macromolecular statistical ensemble of projections obtained from single-particle cryo-EM experiments. Specifically, the following features are incorporated in the simulation: (1) two independent, easily identifiable conformational motions; (2) an exactly defined distribution of occupancies across all states; (3) uniformly sampled projection directions; (4) additive Gaussian noise with appropriate signal-to-noise ratio (SNR); and, (5) experimentally-accurate microscopy parameters for each image, represented by the contrast transfer function (CTF). Given such absolute control and knowledge of the baseline object’s mechanics, both the accuracy of machine-learning algorithms and appropriateness of maximum-likelihood approaches can be quantified with exactitude, thus providing an assessment of these techniques’ strengths and limitations.

## Methods

Our procedure for the creation of a synthetic continuum dataset is outlined in Figure 1. First, a suitable macromolecule (for which a PDB atomic structure exists) is chosen as a foundational model. The atomic coordinates from this PDB file allow the continuum model to be generated at the ground level. Using this atomic structure as a seed, a sequence of additional states is then generated by altering specific subregions of the macromolecule’s structure so as to represent quasi-continuous conformational motions. In this way, many sets of coordinates in PDB format are generated. The number of these mutually decoupled motions defines the number of degrees of freedom, *n*, realized by the system (i.e., the number of independent conformational motions). In sum, this quasi-continuum of states spanning *n* conformational motions defines the molecular machine’s state space. In our example (see Results), *n* = 2 was chosen to demonstrate the simulation procedure.

Every structure created within this state space is then aligned along a common set of 3D coordinates (representing a particular reference structure) to ensure all motions are centered. These centered structures are then converted into 3D Coulomb potential maps, whereby values of electric potential are stored in each voxel. The values in these voxels thus represent the net electrostatic contributions in each region of space from all atomic species within the corresponding PDB-formatted structure file.

At the same time, an occupancy map is created to represent the preference of a molecule to occupy its state, defining the energetics of the state space (i.e., the likelihood of each state to be observed in the experiment). This map is then used to create clones of the 3D Coulomb potential maps. For each 3D Coulomb potential map, an alignment parameter text file is created to store all relevant information for calculating multiple 2D projections of that map. These files contain all of the angular information and experimentally-relevant electron microscopy parameters required to define the characteristics of each 2D projection taken, including modification by the contrast transfer function.

**Figure 1:**
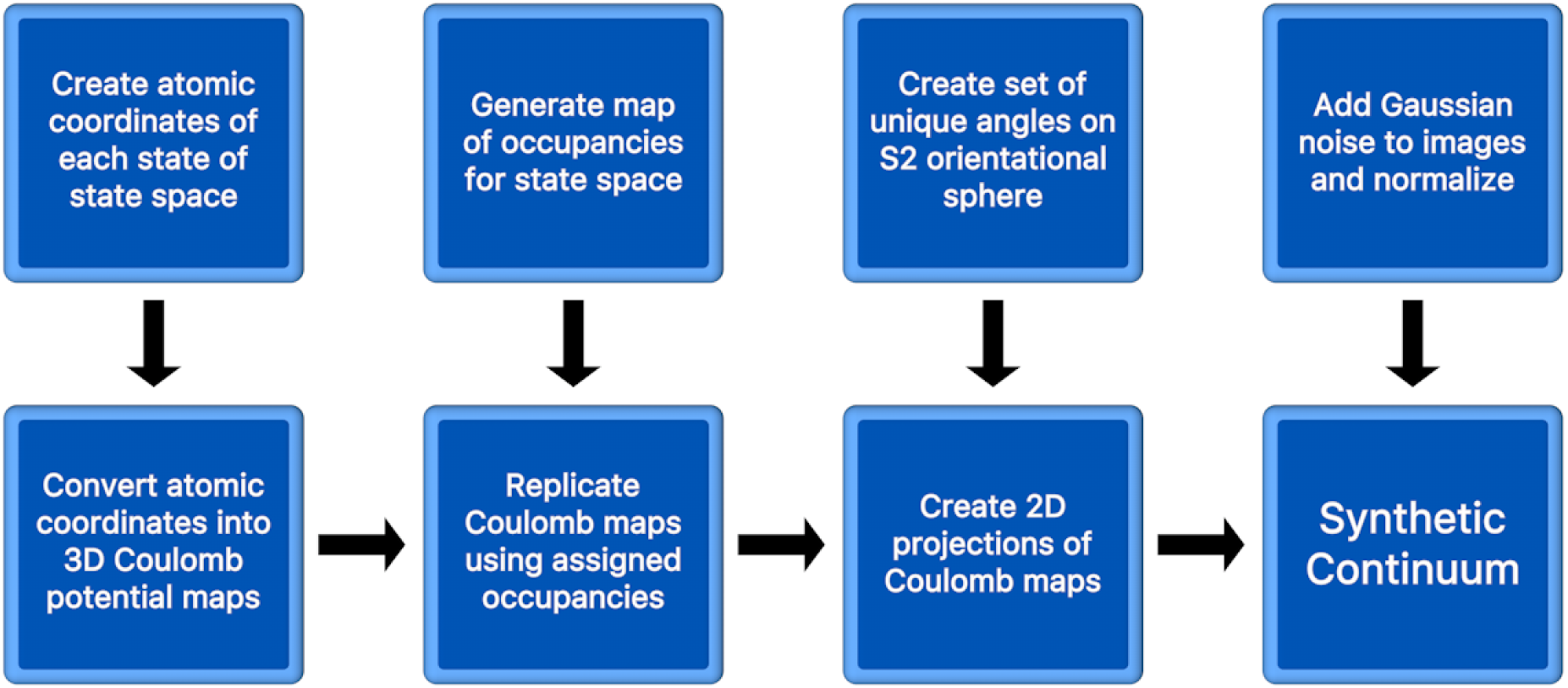
Flowchart representation of the workflow used for creation of the synthetic continuum dataset.

These alignment parameter files are then used as inputs into a 2D projection algorithm. Projections of each alignment file’s corresponding Coulomb potential map are obtained via standard line integrals in selected directions as to mimic projections generated in the transmission electron microscope operated in the bright-field mode. As a result, an image stack of synthetic 2D cryo-EM projections is produced for every 3D Coulomb potential map. Noise is then added to each image across all of the image stacks individually. The final output is a single stack of simulated cryo-EM 2D projections with a corresponding alignment file detailing information for each individual image in that stack. These files in total encompass the entirety of the 3D synthetic continuum as in an idealized cryo-EM experiment.

## Results

The heat shock protein Hsp90^23^ was chosen as a starting structure for this workflow due to its simple design – exhibiting two arm-like subunits (chain A and B, containing 677 residues each) connected together in an overarching V-shape. *In vivo*, these arms are known to close after binding with ATP – with Hsp90 acting as a chaperone to stabilize the structures of surrounding heat-vulnerable proteins. During its work cycle, Hsp90 naturally undergoes large conformational changes, transitioning from its two arms spread open in a full V-shape (inactive state) to both arms bound together along the protein’s central line of two-fold symmetry (active state) following ATP binding. We initiated our workflow with the fully closed state (via entry PDB 2cg9), whose structure was determined at 3.1 Å using X-ray crystallography.^24^

Casting Hsp90’s biological context aside, liberties were taken in the choice of the synthetic model’s leading degrees of freedom. Instead of one singular range of conformational motions (arms open to closed, as *in vivo*), we decided on two unique motions that covered similar distance scales while being both easily identifiable and visually separable. These two conformational motions were designed so as to be fully decoupled from each other (i.e., the number of degrees of freedom *n* = 2), such that no overlap occurred between the two sets of distinct atomic displacements (Figure 2). To further avoid any unintentional coupling between these motions, no minimization of electrostatic energies between all atoms (to correct geometry) in each conformational state was performed. To note, although we believe that two conformational motions meet the minimum level of sophistication required to test machine-learning algorithms, any number of decoupled motions could be defined during this process. As an example of recent use, the manifold embedding technique implemented in ManifoldEM^15^ currently analyzes energetics in sets of a maximum of two conformational motions (as currently restricted by the lack of available datasets with statistically relevant higher-order eigenfunctions).

Atomic manipulations of the original PDB coordinate file were done using PyMOL.^25^ For the first motion (termed Conformational Motion 1, or CM1), the chain A arm was rotated outwards (directly away from chain B) from its central hinge (at residue 677), in 1° increments until a series of 20 rotational states had been defined (Figure 3A). For each of these 20 CM1 states, all residues above chain B’s elbow (residues 12-442) were then rotated perpendicularly to the CM1 motions in 2° increments until a series of 20 rotational states had been defined for the second motion (termed CM2, Figure 3B). These operations resulted in the creation of a total of 400 unique conformational states. The ensemble of these states can be organized in a 20*×*20 state space, whereby each entry represents one of the possible combinations of CM1 and CM2. This state space represents our synthetic model’s complete ensemble of physically allowable conformations (i.e., the quasi-continuum). The specific size of the state space (400 states) was chosen based on the relative scale of the protein and range of its motions, providing approximately 3 Å and 2 Å gaps between states over a total arc length of 60 Å and 40 Å for CM1 and CM2, respectively.

**Figure 2:**
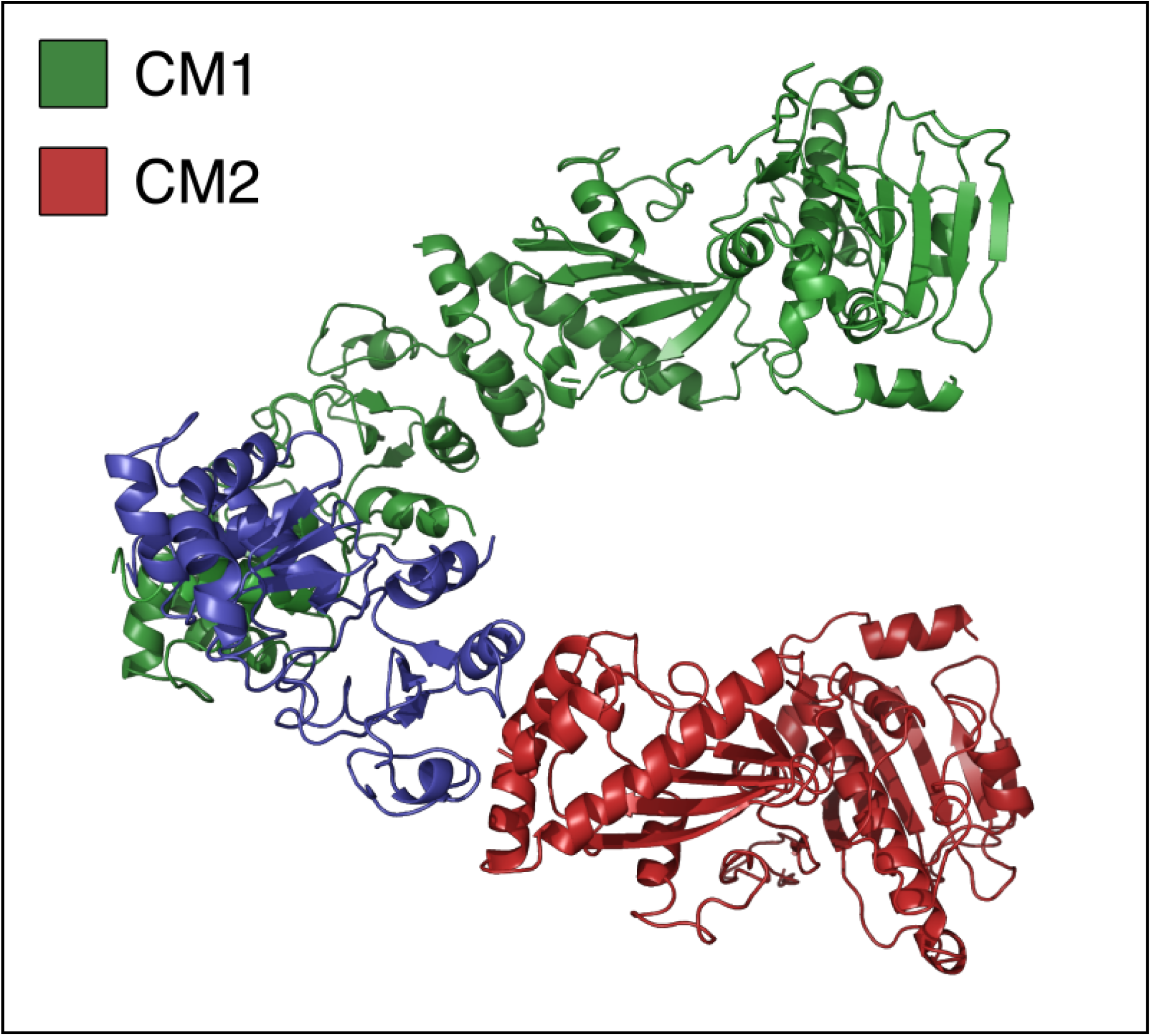
PDB of state 20_01 with residues demarcated for CM1 (green) and CM2 (red). The CM1 central hinge can be found near the intersection of the blue and green regions at residue 674, while the CM2 hinge is found near the intersection of the blue and red regions at residue 443. To note, the residues making up the blue region are immobile throughout the entire state space. As well, the maximum diameter of the protein was measured in this state at approximately 120 Å.

**Figure 3:**
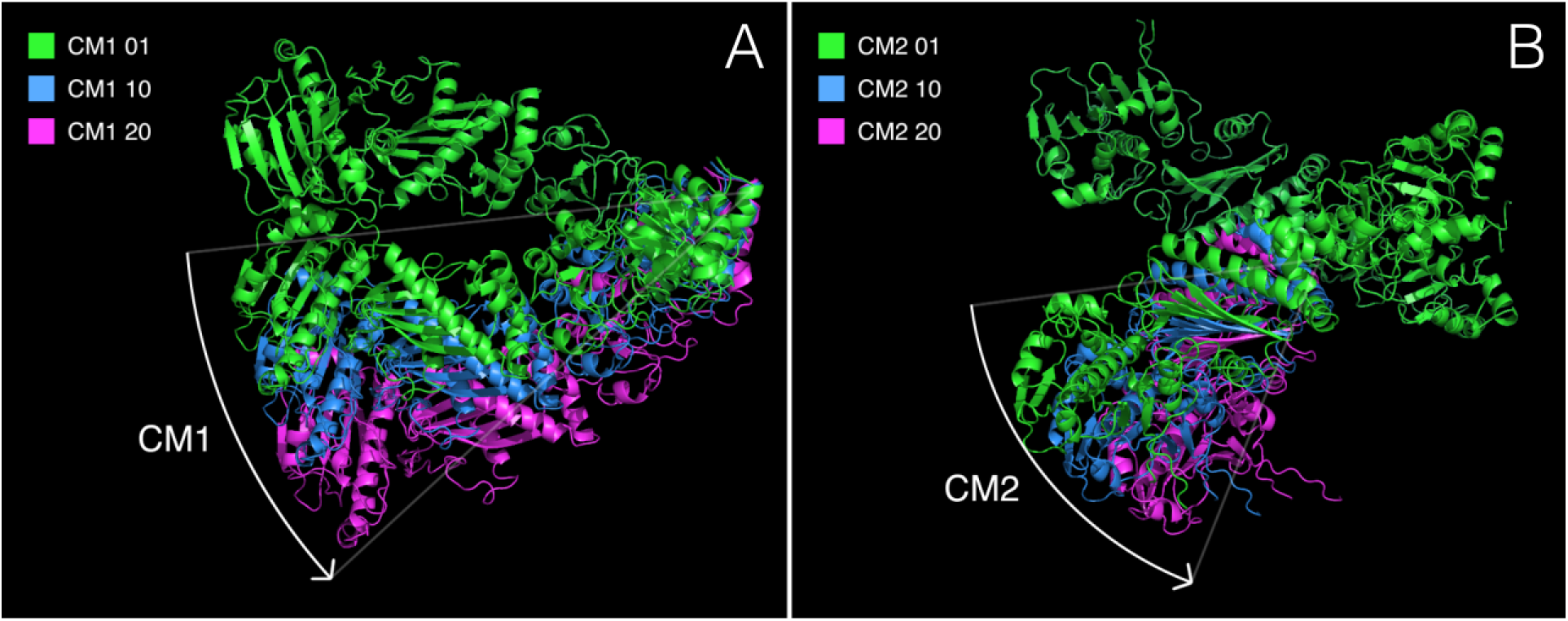
Figure 3A presents a comparison of CM1’s first, middle, and final states (state 01_01, 10_01, and 20_01, respectively). In Figure 3B, a comparison of CM2’s first, middle, and final states are shown (state 20_01, 20_10, and 20_20, respectively). This rotation was performed perpendicularly to CM1’s hinge-rotation, whereby only those atoms above chain B’s elbow-region were perturbed. As a note, to remove the potential for overlapping residues within certain states, residues 1-11 (making up a loose tail) were removed from both chain A and B.

These 400 structures represented by PDB files were then individually transformed into 400 3D Coulomb potential maps (formatted as MRC volume files) with a sampling rate of 1 Å per voxel and simulated resolution of 3 Å (chosen based on the resolution determined for PDB 2cg9) using the EMAN module `e2pdb2mrc`^26^ (Figure 4). As a note, all 3D atomic coordinates were moved to a common frame of reference using Phenix^27^ (as defined by state_01_01’s center of mass) to ensure all motions were centered.

Next, an arbitrarily defined 2D occupancy map was next created and assigned across all members in the state space (Figure 5A) so as to simulate the number of sightings of each conformation that would be obtained via cryo-EM. Each entry in this occupancy distribution is an integer *∈* [1,..,9] used to define the number of clones produced for each state. As the number of sightings of each conformation observed in an experiment directly relates to its underlying energetics, this distribution was then transformed via the Boltzmann factor into its corresponding energy landscape (Figure 5B).

A unique RELION^29^ alignment parameter file (in STAR format) was created for each of the 1000 Coulomb potential maps (consisting of the original 400 maps plus their multiple-occupancy clones). The STAR format of RELION was used for organizing and storing information on each of the 2D snapshots obtained from the simulated cryo-EM experiments. By convention, each row in the alignment file represents a single snapshot of a macromolecule from the experimental ensemble, with its columns detailing that snapshot’s corresponding microscopy parameters. As each experimentally-obtained image naturally captures a macro-molecule in a range of different spatial orientations, angular assignments were given to map the position of each image within the orientational space. The location of each snapshot in this space is represented by a combination of Euler angles, which are provided in the columns for each row within the alignment file. Along with this information, additional columns carry the location of the macromolecule’s center of mass in the image, the defocus values of the image, and other relevant parameters such as voltage, spherical aberration, and amplitude contrast ratio.

**Figure 4:**
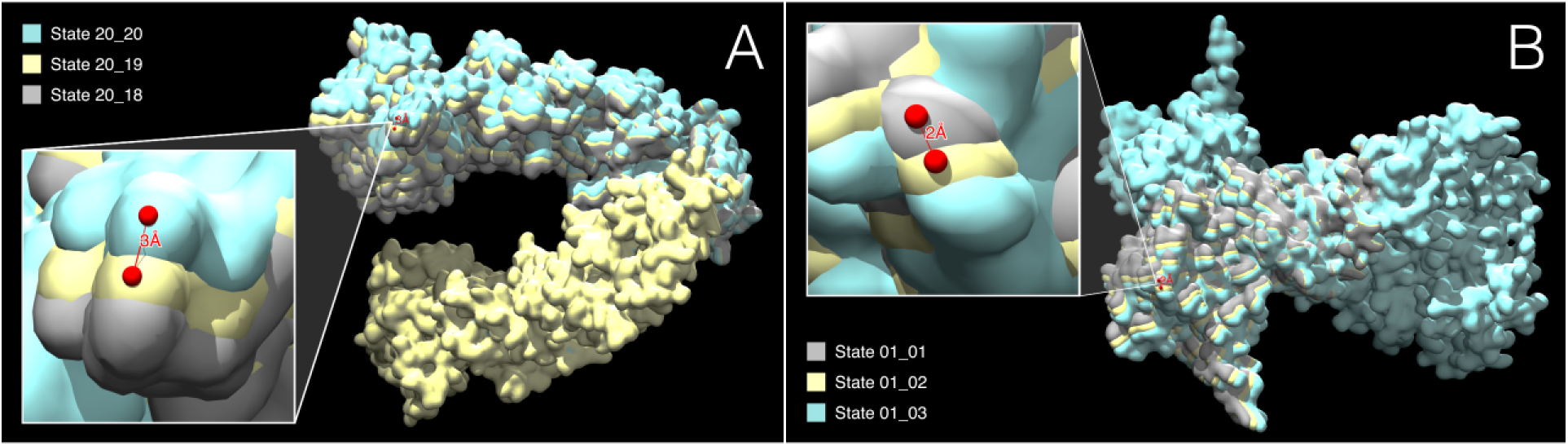
Figure 4A presents a volumetric overlay of first three rotational states of CM1 (state 01_01, state 02_01, state 03_01), visualized as Coulomb potential maps (MRC format) via Chimera.^28^ As can be seen, only the upper arm (chain A) has been rotationally displaced along CM1, with 3 Å gaps measured between each consecutive state at the peripheral ends of this rotated region. Figure 4B presents a volumetric overlay of last three rotational states of CM2 (state 20_18, state 20_19, state 20_20). Only the upper region of chain B (above the elbow) has been rotationally displaced along CM2, with 2 Å gaps measured between each consecutive state at the peripheral ends of this rotated region. To ensure that the two conformational motions (CM1 and CM2) were completely decoupled, no interpolation techniques (such as implemented in PyMOL’s “morph” command) were used for state generation – as these morphs propagate small movements between atoms that could weakly couple CM1 with CM2.

To create simulated snapshots of Hsp90 within this framework, we needed to first create alignment files that contained the relevant parameters needed for subsequent 2D projection operations to be performed. Given this agenda, we created an alignment file with 812 rows for each volume in our synthetic dataset. These 812 rows were each given a set of Euler angles from a uniformly spread distribution in orientational space, representing 812 vertices (viewing directions of that state of the molecular machine) on an evenly tessellated sphere (Figure 6). For each row within every alignment file, a value for contrast transfer function (CTF) parameters was randomly assigned from within an experimentally relevant range (defocus values assigned evenly between 2,000 and 10,000 Å). Constant values were used for voltage (300 kV), spherical aberration coefficient (2.7 mm), and amplitude contrast ratio (0.1) to emulate typical electron microscopy parameters. Positional *x* and *y* shifts were set to zero so as to avoid application of image padding artifacts during re-centering (a problem that does not exist for experimental data taken from micrographs). Ultimately, each of these 812 rows represents the collection of information required to form a unique, virtually accurate cryo-EM snapshot as taken from a given orientational direction of its corresponding Coulomb potential map. In total, the rows of these alignment files provide the instructions for creating 812,000 (812*×*1000) snapshots taken across all combinations of orientations and conformations throughout our entire ensemble.

**Figure 5:**
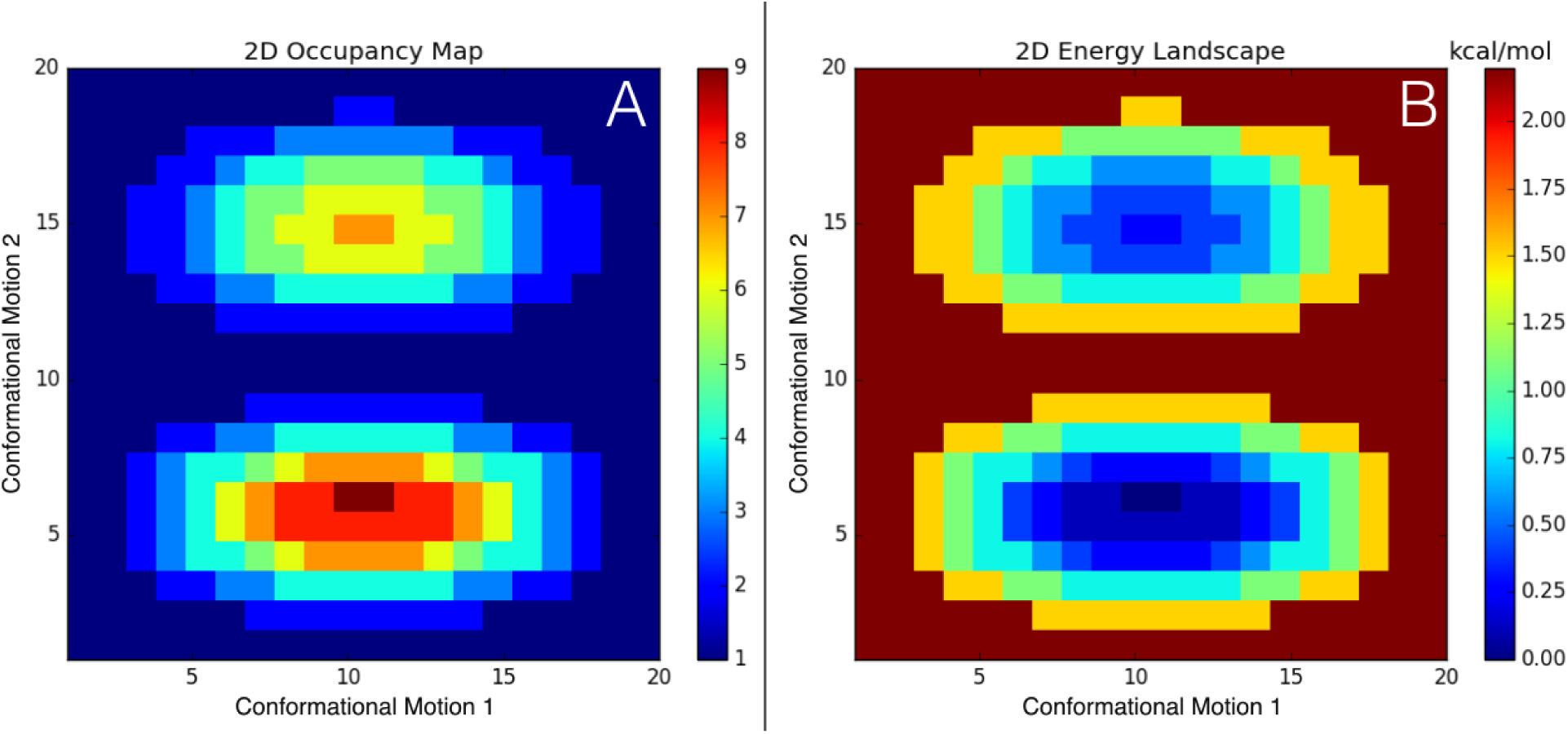
Figure 5A presents the 2D distribution of occupancies for all 400 states in the state space. The formula^1^ for this occupancy map was chosen as to provide easily distinguished features along both 1D (Figure 9) and 2D conformational motions. As an example of use, only the original copy of state 01_01’s Coulombic potential map was used, while 9 copies of state 10_06’s Coulombic potential map were created for use in all subsequent steps. In total, the creation of these clones increased the number of MRC volume files was increased from 400 to 1000. Figure 5B presents the transformation of the occupancy map (left) into its corresponding energy landscape via the Boltzmann factor: −log(m_i_*/*m), where m_i_ is the occupancy of the current state and m is the maximum occupancy across all states. Within this distribution, the state with the lowest energy is defined at zero kcal/mol, while the highest energy state has an energy of 2.2 kcal/mol (occurring at states 6_10 and 6_11).

**Figure 6:**
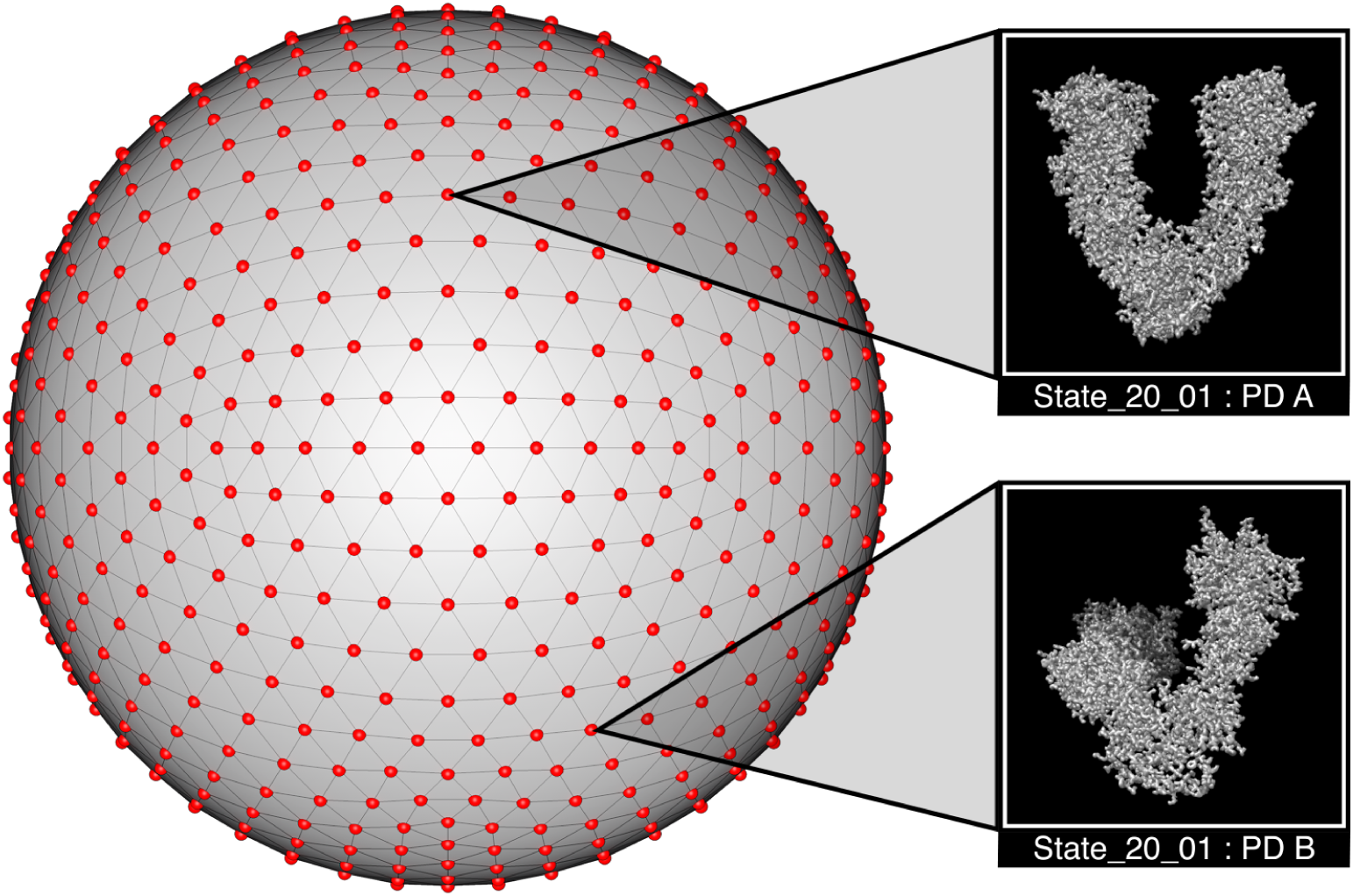
Distribution of projection directions (PDs, shown as red points) as arranged across the space of all possible viewing-angles (S2 sphere). This tessellation was achieved by forming an icosahedron with 48 segments in Cinema4D^30^ to create 812 evenly spaced vertices (with edges seen in black) representing a set of available PDs. Next, the Cartesian coordinates of each of these vertices were transformed into spherical coordinates with in-plane rotation set to zero. Given this tessellation, each vertex (PD) is approximately 6° from its nearest neighbor. A quaternion transformation was next performed on each of these coordinates (see Supplementary Material) to emulate small, random angular perturbations about each PD. Importantly, these perturbations are designed to reproduce the naturally-occurring orientational variations of a real experiment.

We next provided this collection of information as inputs into RELION’s 2D projection module (`relion_project`). This operation created a 3D Fourier transform of each volume from which 812 2D central sections were selected (with interpolation) and multiplied with the corresponding CTF before being transformed back into real space. ^29^ In this way, a unique 2D projection was obtained for each combination of Euler angles in the corresponding alignment file row. In all, this procedure resulted in the creation of 812 unique 2D images for each of our 1000 volumes, with this information stored in 1000 image stacks (MRC format) (Figure 10).

Next, all 1000 image stacks (i.e., one from each volume) were combined into one single image stack while adding Gaussian noise to each image individually such that its signal-to-noise ratio (SNR) became 0.1. This value has been previously established as a suitable approximation for experimental SNR in images obtained by cryo-EM, and can be attributed to the low contrast between macromolecules and their surrounding ice, as well as the limited electron dose required to avoid radiation damage. ^31,32^ We define this ratio by the division of the image’s variance in signal 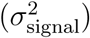 by its variance in noise 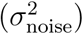^4^ Here, *signal* represents the 2D region of pixels corresponding to the average area occupied by the protein, obtained by masking out all pixels within 0.5 standard deviations of each image’s mean intensity value corresponding to the approximately uniform-intensity background.

This process was thus performed by first finding the mean pixel intensity (*µ*_signal_) and variance 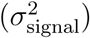 of the *signal*, and calculating 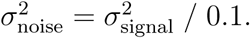 Using these statistics, we then apply additive Gaussian noise to each image in order to obtain an output image having the desired SNR. During this process, a sample from the Gaussian distribution, generated with NumPy^33^ via `numpy.random.normal`(*µ*_signal_, *σ*_noise_, (`rows, columns`)), is added to each pixel’s intensity. Each resulting image was then normalized such that the average pixel intensity and standard deviation of pixel intensities was approximately 0 and 1, respectively. Next, `mrcfile`^34^ was used to combine all images into a single image stack (Figure 7). Finally, a single alignment file was produced containing the original parameters for each of the 812,000 images within the image stack.

**Figure 7:**
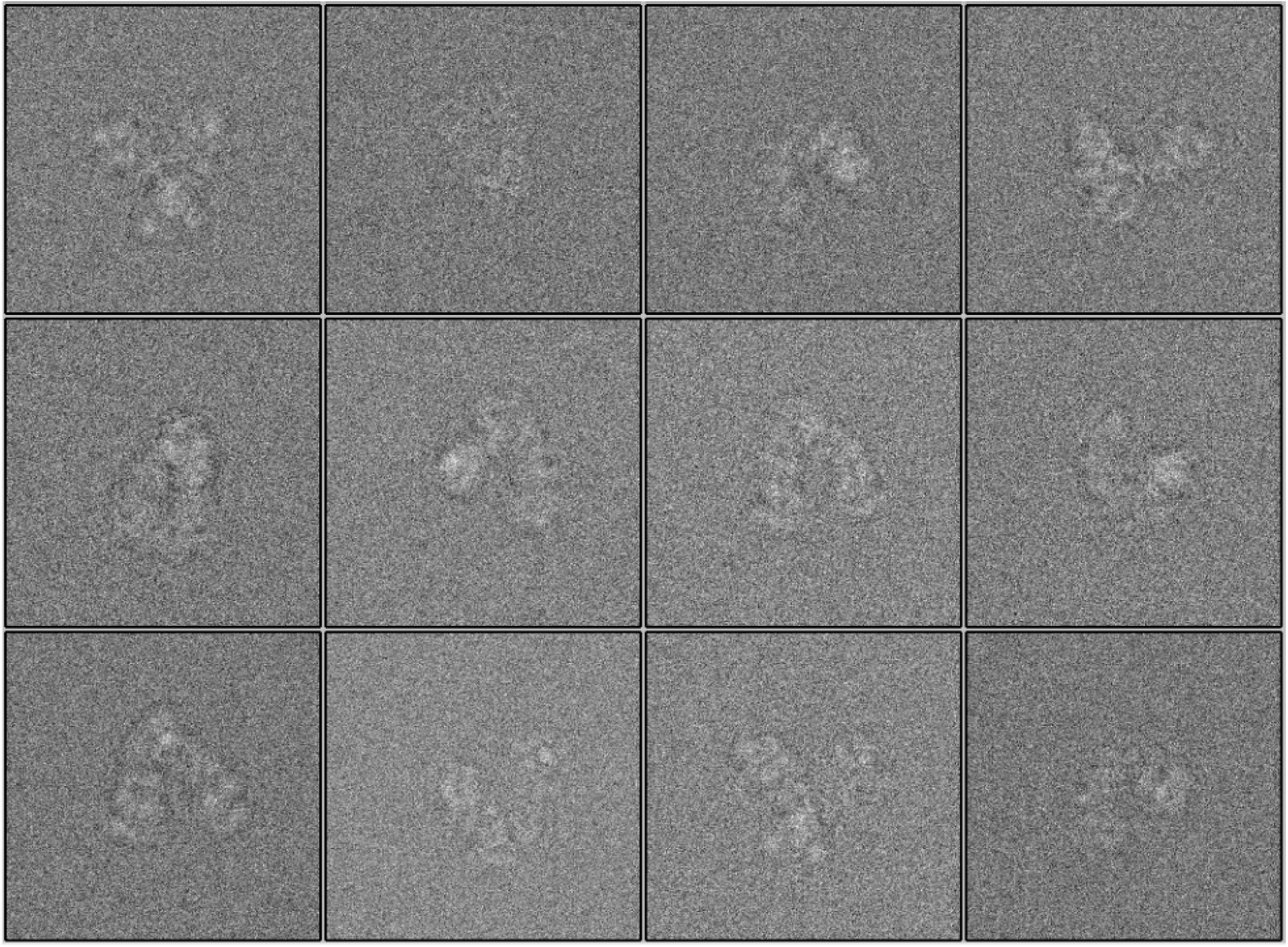
Small selection of images as obtained from the tail end of the final MRCS image stack (within the region of state 20_20). Each image was uniquely normalized after applying additive Gaussian noise with SNR = 0.1.

## Conclusion

An ensemble of projections of the Hsp90 protein undergoing quasi-continuous conformational changes has been provided as an exemplary ground-truth model for exploring the accuracy and appropriateness of techniques claiming to specialize in the reconstruction of structure and function from cryo-EM data. Along with this publicly-available dataset, we have provided a workflow for generating cryo-EM simulated data from atomic models subjected to user-defined conformational changes. This workflow can be completely customized to meet the needs of the user, including choice of atomic model, number and types of conformational motions, density and distribution of states, energetics of state space, distribution of projection directions, microscopy parameters and noise. This flexibility will facilitate a comprehensive investigation of external data-analytic techniques, highlighting the performance and limitations of these approaches under a number of complex conditions for which the user has absolute control. All scripts for reproducing and customizing this workflow have been made available online.^1^

## Acknowledgement

We thank Abbas Ourmazd, Ali Dashti, and Ghoncheh Mashayekhi for thoughtful feedback on ManifoldEM results across a range of synthetic prototypes, and Suvrajit Maji and Hstau Liao for helpful comments, insights and review throughout development. This research was supported by the National Institutions of Health Grants GM29169 and GM55440 (to J.F.).

## Supporting Information Available

**Figure 8:**
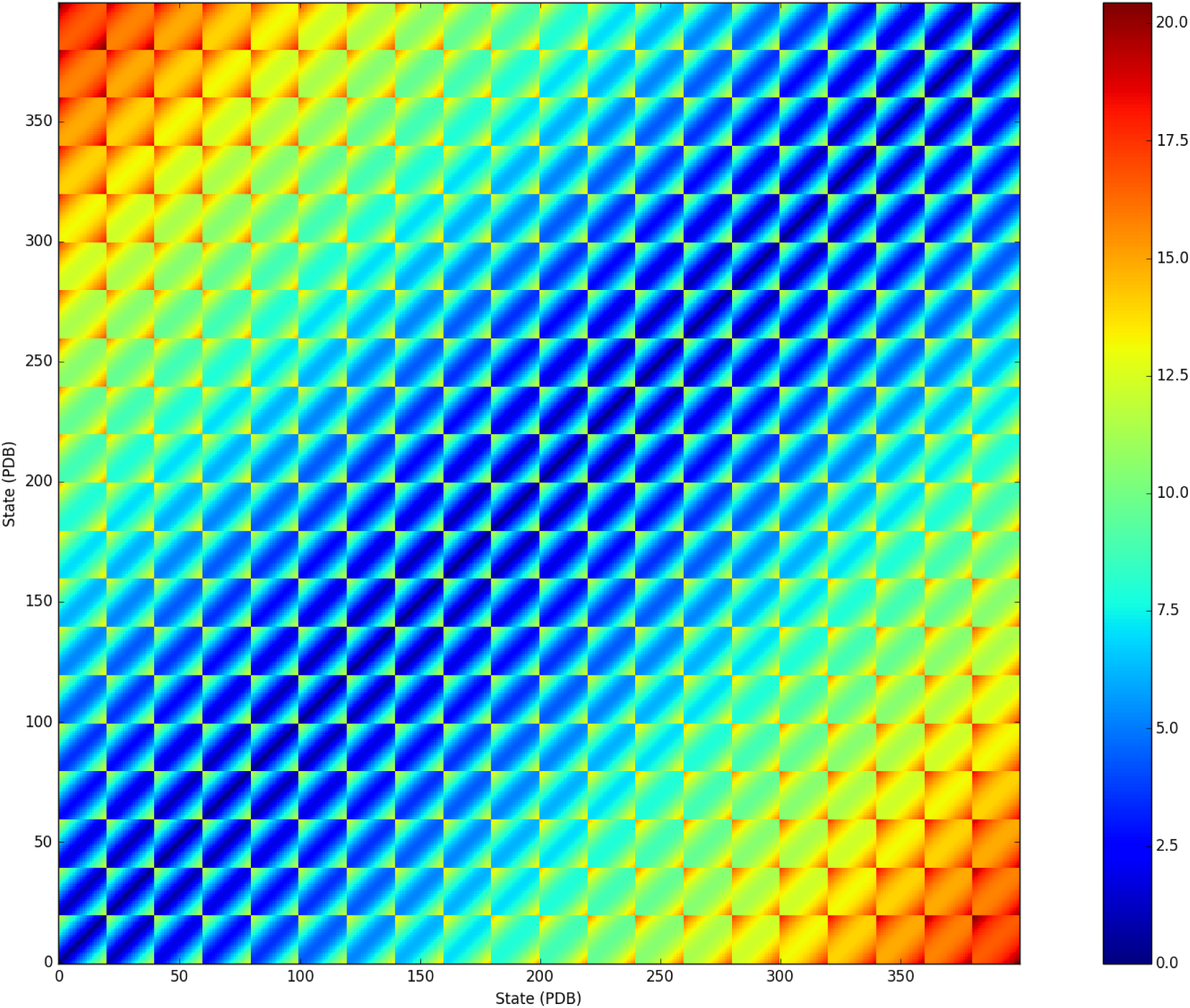
RMSD (via PyMOL’s ‘rms_cur’ command) of all combinations of atomic structures (PDBs) in the state space. The indices (1, 2, …, 400) represent a sequential ordering along the columns in the 20×20 CM1-CM2 state space, whereby index 1 corresponds to state 01_01, index 2 to state 01_02, and index 21 to state 02_01. Further, the entries in this 400×400 RMSD matrix provide a quantitative measure of similarity between every pair of 3D protein structures – effectively representing the ground-truth distance matrix for the continuum. Two unique patterns consisting of diagonal-pinstripes and square-boxes can be readily identified within the matrix and correspond to CM1 and CM2, respectively. The smallest distance between structures is 0.64 Å and occurs between adjacent CM2 states (e.g., the RMSD between state 01_01 and state 01_02). The largest distance is 20.44 Å and occurs between state 01_01 and state 20_20.

**Figure 9:**
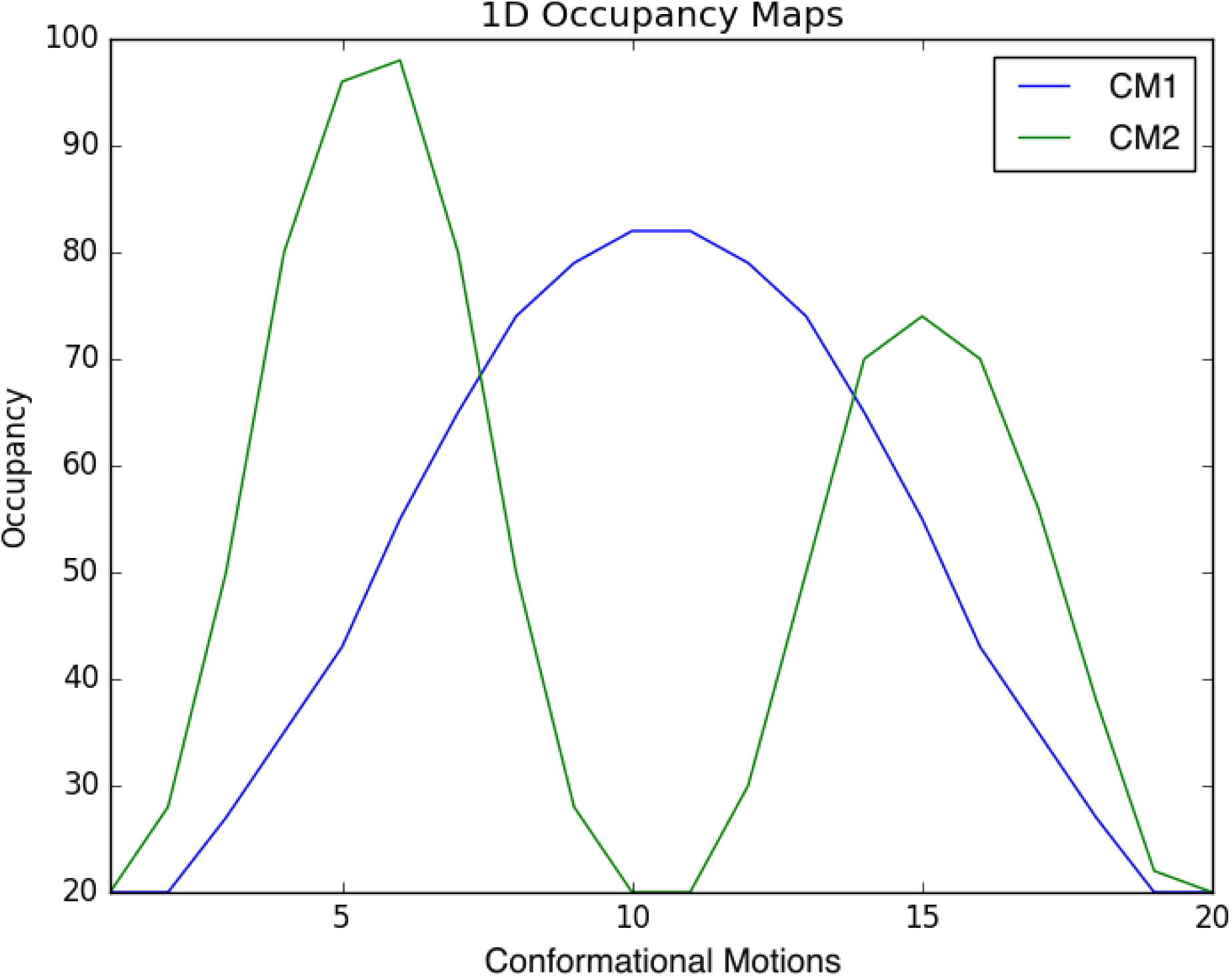
1D distribution of occupancies for all 400 states as projected onto CM1 and CM2. This occupancy distribution has been formulated such that in 1D, both conformational motions are easily distinguishable. Namely, CM1 and CM2 have a modal and bimodal distribution, respectively, whereby CM2 has also been given a wider range of occupancy values.

**Figure 10:**
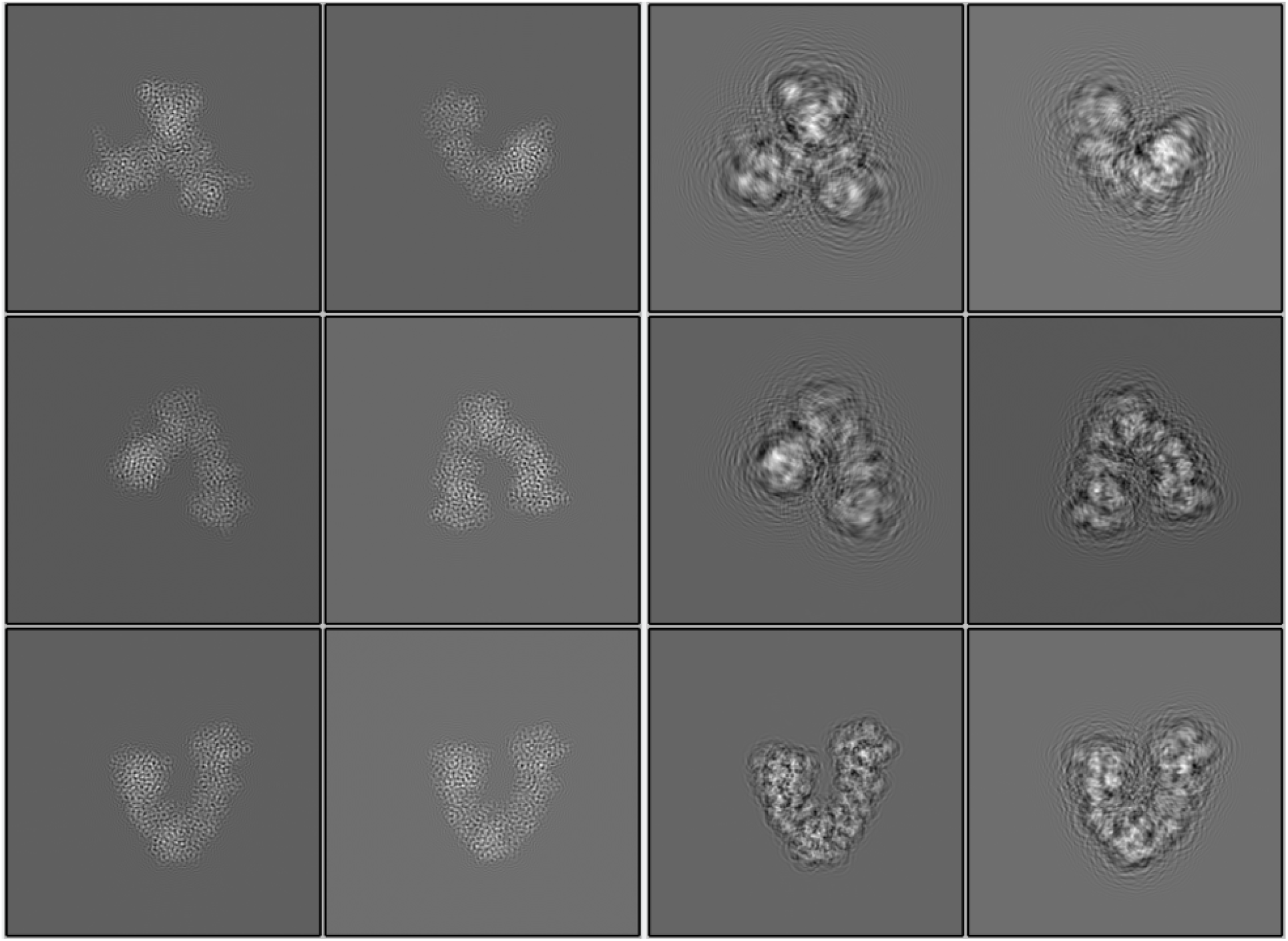
Comparison of state 20_20’s image stack with and without applied defocus values (left and right side, respectively) using RELION’s “project” command. Note that the combination of defocus values range and window size (250 pixels) was chosen so as not to lose information scattered due to the ripples of the point spread function into the regions outside of each image’s borders.

### PD Rotations

Each small, random angular-perturbation about each image’s corresponding PD was performed via two sequential quaternion transformations. These transformations were designed to preserve uniform rates of angular changes for all images across the S2 sphere (Figure 11A). First, the initial combination of Euler angles for a given image’s corresponding PD (Figure 6) was transformed into an equivalent unit vector 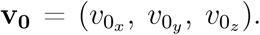 Next, a unit vector, **w** = (*w*_*x*_, *w*_*y*_, *w*_*z*_), orthogonal to **v**_0_, was generated. A unit quaternion was generated to represent a rotation about the axis **w** by an angle *θ*_1_ as follows:

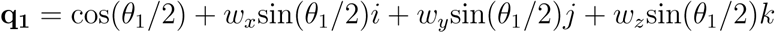

The rotation angle *θ*_1_ for each image was chosen from a random Gaussian distribution with zero mean (center of the PD) and a standard deviation of 1.25°. The vector **v_0_** was then rotated according to **v_1_** = **q_1_** ⋅ **v_0_** ⋅ **q_1_**^*−*1^, which effectively rotated the vector **v_0_** radially outward from its original location (PD center) on a great circle, spanning a range of approximately 6° on the S2 sphere. Next, **v_1_** was randomly rotated by a second unit quaternion with axis of the PD’s original location, **v_0_**, by *θ*_2_.

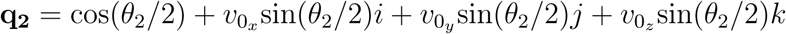

The angle *θ*_2_ was chosen randomly from a uniform distribution between *−*180° and 180° for each image, such that the initial Gaussian distribution across each PD (formed from numerous possible random samples of **v_1_** locations for each image) was transformed into an isotropic Gaussian distribution about the center of each PD (Figure 12). This second transformation, **v_2_** = **q_2_** ⋅ **v_1_** ⋅ **q_2_**^*−*1^, defined the final vector for each image, which was subsequently converted into its corresponding Euler angle representation.

**Figure 11:**
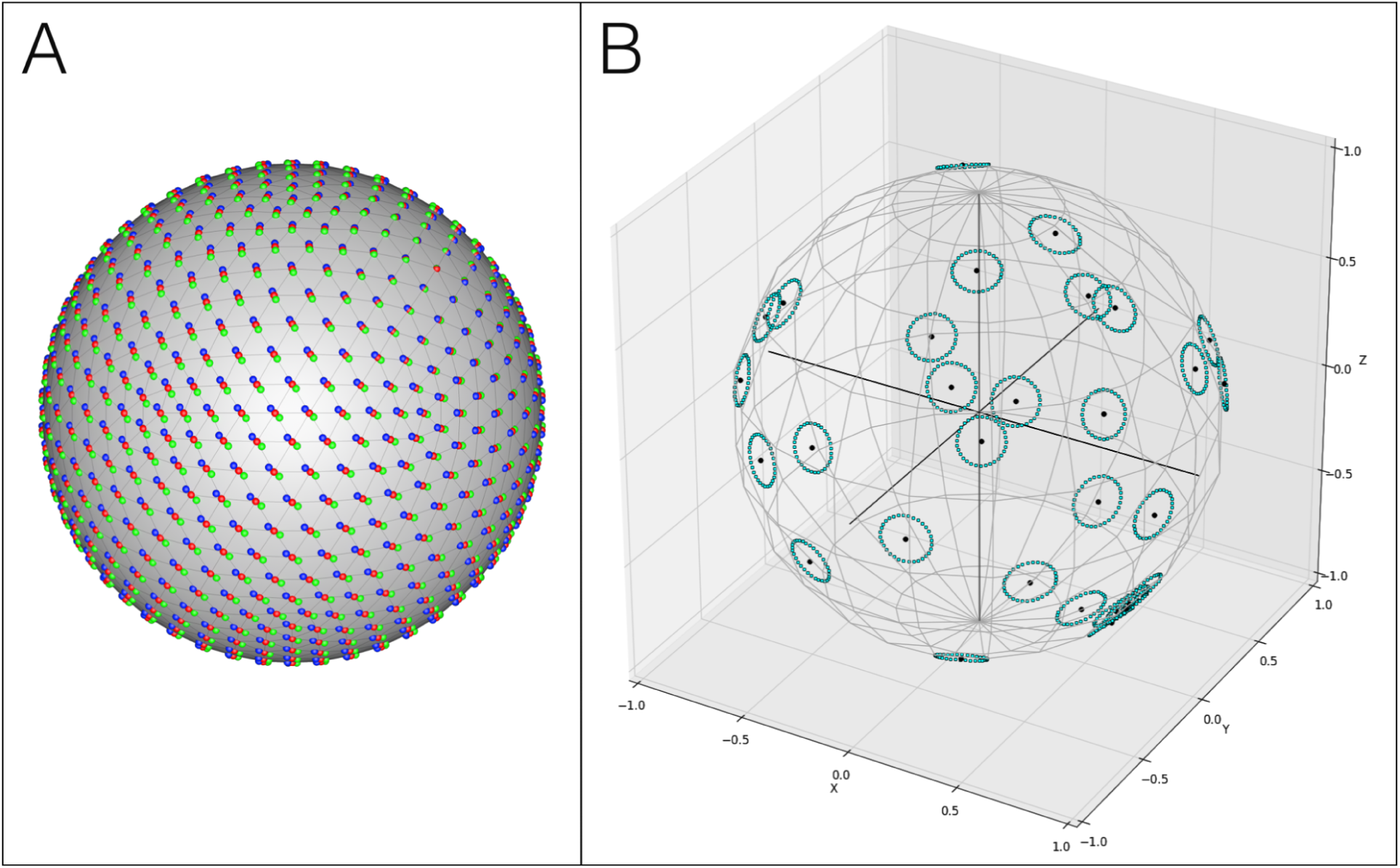
Figure 11A schematizes the problem of applying global rotations to all PDs, which results in a non-uniform angular distribution on the S2 sphere. The original PD locations are represented by red vertices, while two oppositely-directed global rotations are shown as blue and green vertices. As can be seen, the distribution of changes of blue and green vertices about each corresponding red vertex are non-uniform, and vanishes near the poles formed by the axis of rotation. Figure 11B presents a schematic of the effects of the back-to-back quaternion transformations, resulting in a uniform distribution on the entire S2 sphere. In this example, the first quaternion transformation rotates each point radially away from its PD center by an equal amount 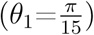 while the second quaternion transformation incrementally rotates those points 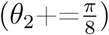 along a circle enclosing the original PD with angular radius *θ*_1_.

**Figure 12:**
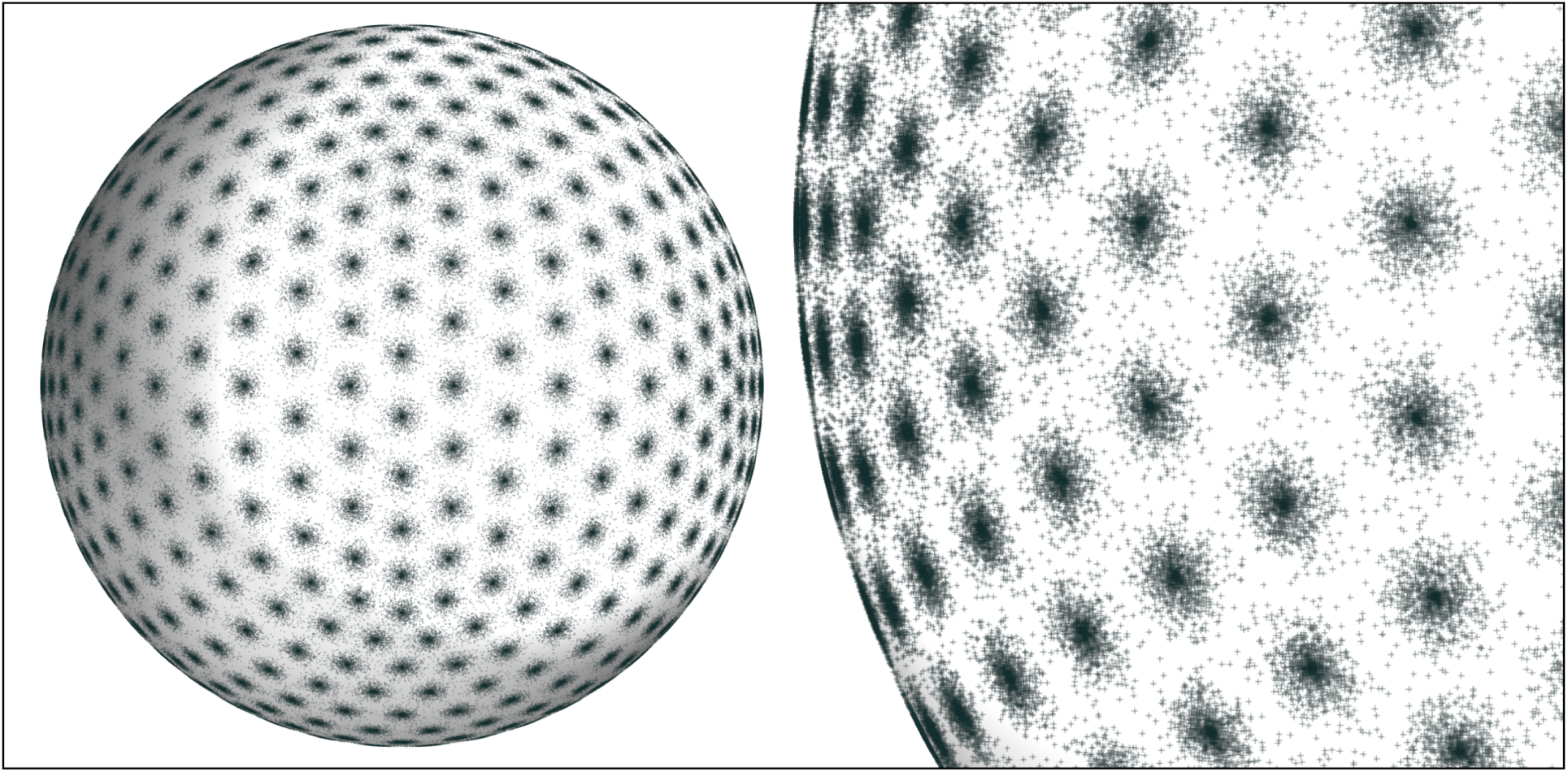
The final distribution of all 812,000 images on the S2 sphere after the back-to-back quaternion transformations with randomized sampling of *θ*_1_ and *θ*_2_ for each image. As can be seen, the Gaussian distribution defining the radial distance of the first transformation has been chosen such that little overlap occurs between the locations of images in neighboring PDs. This ensures that the energy landscape within each PD is locally conserved, such that both global-manifold and multiple-manifold (per PD) approaches can directly compare their algorithm’s analysis to the ground-truth energetics.

## Notes

https://github.com/evanseitz/cryoEM_synthetic_continua

## References

(1) Seitz, E. Repository: Cryo-EM Synthetic Continua (Initial Release). Zenodo 2019, 10.5281/zenodo.3561105.

(2) Frank, J. Generalized single-particle cryo-EM: a historical perspective. Microscopy 2016, 65, 3–8.

(3) Frank, J. Single-Particle Reconstruction – Story in a Sample. Nobel Lecture 2017,

(4) Frank, J. Three-Dimensional Electron Microscopy of Macromolecular Assemblies: Visualization of Biological Molecules in Their Native State; Oxford University Press: Oxford, New York, 2006.

(5) Whitford, P. C.; Altman, R. B.; Geggier, P.; Terry, D. S.; Munro, J. B.; Onuchic, J. N.; Spahn, C. M. T.; Sanbonmatsu, K. Y.; Blanchard, S. C. Dynamic views of ribosome function: Energy landscapes and ensembles; Springer, Vienna, 2011.

(6) Liu, W.; Boisset, N.; Frank, J. Estimation of variance distribution in three-dimensional reconstruction. II. Applications. Journal of the Optical Society of America. A, Optics, image science, and vision 1996, 12, 2628–35.

(7) Penczek, P. Variance in three-dimensional reconstructions from projections. 2002, 749–752.

(8) Penczek, P.; Yang, C.; Frank, J.; Spahn, C. Estimation of variance in single-particle reconstruction using the bootstrap technique: The Path Toward Atomic Resolution/ Selected Papers of Joachim Frank with Commentaries. 2018, 389–404.

(9) Penczek, P.; Frank, J.; Spahn, C. A method of focused classification, based on the bootstrap 3D variance analysis, and its application to EF-G-dependent translocation. Journal of structural biology 2006, 154, 184–94.

(10) Penczek, P.; Kimmel, M.; Spahn, C. Identifying Conformational States of Macromolecules by Eigen-Analysis of Resampled Cryo-EM Images. Structure (London, England: 1993) 2011, 19, 1582–90.

(11) Liao, H.; Frank, J. Classification by bootstrapping in single particle methods. Proceedings / IEEE International Symposium on Biomedical Imaging: from nano to macro. IEEE International Symposium on Biomedical Imaging 2010, 2010, 169–172.

(12) Dashti, A.; Schwander, P.; Langlois, R.; Fung, R.; Li, W.; Hosseinizadeh, A.; Liao, H. Y.; Pallesen, J.; Sharma, G.; Stupina, V. A.; Simon, A. E.; Dinman, J. D.; Frank, J.; Ourmazd, A. Trajectories of the ribosome as a Brownian nanomachine. Proceedings of the National Academy of Sciences 2014, 111, 17492–17497.

(13) Schwander, P.; Fung, R.; Ourmazd, A. Conformations of macromolecules and their complexes from heterogeneous datasets. Philosophical Transactions of the Royal Society B 2014,

(14) Frank, J.; Ourmazd, A. Continuous Changes in Structure Mapped by Manifold Embedding of Single-Particle Data in Cryo-EM. Methods 2016, 100.

(15) Dashti, A.; Hail, B. D.; Mashayekhi, G.; Schwander, P.; des Georges, A.; Frank, J.; Ourmazd, A. Functional Pathways of Biomolecules Retrieved from Single-particle Snapshots. 2018.

(16) Coifman, R.; Lafon, S.; Lee, A.; Maggioni, M.; Nadler, B.; Warner, F.; Zucker, S. Geometric diffusions as a tool for harmonic analysis and structure definition of data: Diffusion Maps. Proceedings of the National Academy of Sciences of the United States of America 2005, 102, 7426–31.

(17) Coifman, R.; Lafon, S. Diffusion maps. Applied and Computational Harmonic Analysis 2006, 21, 5–30.

(18) Moscovich, A.; Halevi, A.; andén, J.; Singer, A. Cryo-EM reconstruction of continuous heterogeneity by Laplacian spectral volumes. 2019.

(19) Sorzano, C. et al. Survey of the analysis of continuous conformational variability of biological macromolecules by electron microscopy. Acta Crystallographica Section F Structural Biology Communications 2019, 75.

(20) Zhong, E.; Bepler, T.; Davis, J.; Berger, B. Reconstructing continuously heterogeneous structures from single particle cryo-EM with deep generative models. 2019.

(21) Fischer, N.; Konevega, A.; Wintermeyer, W.; Rodnina, M.; Stark, H. Ribosome dynamics and tRNA movement by time-resolved electron cryomicroscopy. Nature 2010, 466, 329–33.

(22) Ourmazd, A. Cryo-EM, XFELs and the structure conundrum in structural biology. Nature Methods 2019, 16, 941–944.

(23) Goodsell, D. PDB-101 Molecule of the Month: Hsp90. 2008.

(24) Ali, M.; Roe, S.; Vaughan, C.; Meyer, P.; Panaretou, B.; Piper, P.; Prodromou, C.; Pearl, L. Crystal structure of an Hsp90–nucleotide–p23/Sba1 closed chaperone complex. Nature 2006, 440, 1013–1017.

(25) Schrödinger, L. The PyMOL Molecular Graphics System. 2015.

(26) Tang, G.; Peng, L.; Baldwin, P.; Mann, D.; Jiang, W.; Rees, I.; Ludtke, S. EMAN2: An extensible image processing suite for electron microscopy. Journal of structural biology 2007, 157, 38–46.

(27) Adams, P. et al. PHENIX: a comprehensive Python-based system for macromolecular structure solution. Acta Crystallogr D Biol Crystallogr 66(pt 2):213–221. Acta Crystallographica Section D Biological Crystallography 2010, 66, 213-21.

(28) Pettersen, E.; Goddard, T.; Huang, C.; Couch, G.; Greenblatt, D.; Meng, E. UCSF Chimera—A visualization system for exploratory research and analysis. J Comput Chem 2004, 25.

(29) Scheres, S. RELION: Implementation of a Bayesian approach to cryo-EM structure determination. Journal of Structural Biology 2012, 180, 519–530.

(30) GmbH, M. C. Cinema4D. 1985–2019.

(31) Baxter, W.; Grassucci, R.; Gao, H.; Frank, J. Determination of Signal-to-Noise Ratios and Spectral SNRs in Cryo-EM Low-dose Imaging of Molecules. Journal of structural biology 2009, 166, 126–32.

(32) Scheres, S.; Núñez-Ramírez, R.; Gómez-Llorente, Y.; San Martín, C.; Eggermont, P.; Carazo, J. Modeling Experimental Image Formation for Likelihood-Based Classification of Electron Microscopy Data. Structure (London, England: 1993) 2007, 15, 1167–77.

(33) van der Walt, S.; Colbert, S.; Varoquaux, G. The NumPy Array: A Structure for Efficient Numerical Computation. Computing in Science and Engineering 2011, 13, 22–30.

(34) Burnley, T.; Palmer, C.; Winn, M. Recent developments in the CCP-EM software suite. Acta Crystallographica Section D Structural Biology 2017, 73, 469–477.

